# A genome-wide screen links peroxisome regulation with Wnt signaling through RNF146 and tankyrase

**DOI:** 10.1101/2024.02.02.578667

**Authors:** Jonathan T. Vu, Katherine U. Tavasoli, Lori Mandjikian, Connor J. Sheedy, Julien Bacal, Meghan A. Morrissey, Chris D. Richardson, Brooke M. Gardner

## Abstract

Peroxisomes are membrane-bound organelles harboring metabolic enzymes. In humans, peroxisomes are required for normal development, yet the genes regulating peroxisome function remain unclear. We performed a genome-wide CRISPRi screen to identify novel factors involved in peroxisomal homeostasis. We found that inhibition of RNF146, an E3 ligase activated by poly(ADP-ribose), reduced the import of proteins into peroxisomes. RNF146-mediated loss of peroxisome import depended on the stabilization and activity of the poly(ADP-ribose) polymerase tankyrase, which binds the peroxisomal membrane protein PEX14. We propose that RNF146 and tankyrase regulate peroxisome import efficiency by PARsylation of proteins at the peroxisome membrane. Interestingly, we found that the loss of peroxisomes increased tankyrase and RNF146-dependent degradation of non-peroxisomal substrates, including the beta-catenin destruction complex component AXIN1, which was sufficient to alter the amplitude of beta-catenin transcription. Together, these observations not only suggest previously undescribed roles for RNF146 in peroxisomal regulation, but also a novel role in bridging peroxisome function with Wnt/beta-catenin signaling during development.

## Introduction

The peroxisome is a membrane-bound organelle that harbors enzymes for specialized metabolic reactions. The most conserved peroxisomal functions include the beta-oxidation of fatty acids and regulation of reactive oxygen species [Wanders and Waterham 2006]; however, cells tune peroxisome function according to need. For example, peroxisomes in the large intestine of mice contain enzymes for optimal plasmalogen synthesis, while peroxisomes in the small intestines contain enzymes for optimal beta-oxidation of fatty acids [Morvay et al 2017]. Peroxisome function differentiates alongside cell type: for example, in inner ear cells, sound-induced autophagy of peroxisomes protects against noise overexposure [Defourny et al 2019], while in macrophages, peroxisomal metabolism improves phagocytosis [Di Cara et al 2017]. Accordingly, mutations in peroxisomal genes in humans cause a spectrum of Peroxisome Biogenesis Disorders (PBDs) with phenotypes ranging in severity from early infant mortality, developmental abnormalities, and liver dysfunction to more specific metabolic syndromes, sensorineural hearing loss, and retinal degeneration [Braverman et al 2016]. It is therefore important to know both the genes dedicated to peroxisome function in human cells, as well as the mechanisms by which peroxisome abundance and function are coordinated to meet the needs of cell.

Peroxisomes are made and maintained by approximately 35 PEX proteins which coordinate the biogenesis of peroxisome membranes and the import of peroxisomal matrix localized enzymes. Protein import into peroxisomes depends on the presence of peroxisome structures, as well as on many of the best conserved PEX proteins that ensure the efficiency of import. Proteins tagged with a C-terminal peroxisomal targeting signal (PTS1) are recognized by the receptor PEX5, which shuttles the PTS1-cargo to the PEX13/PEX14 docking complex for import across the peroxisomal membrane [Dammai et al 2001; Skowyra et al. 2022]. After import, PEX5 is recycled via extraction by PEX1/PEX6/PEX26 from the peroxisomal membrane following ubiquitination by the PEX2/PEX10/PEX12 E3 ligase complex [Platta et al 2009; Platta et al 2005]. Cells fine tune peroxisomal protein import, and therefore peroxisome function, according to need. The repertoire of imported enzymes is regulated through transcription, as well as ribosomal readthrough that can create protein isoforms with an appended PTS1 tag [Stiebler et al 2014]. The efficiency of import is also regulated cell-wide, for example, phosphorylation of PEX5 by ATM, a DNA repair kinase, can induce peroxisome-specific autophagy in response to oxidative stress [Zhang et al 2015]. Thus, peroxisome homeostasis is tightly regulated in cells and disruption of this regulation can have severe consequences on organismal development. However, the full regulatory network that governs the steady state equilibrium of peroxisome abundance, function, and homeostasis in human cells remains elusive.

Here we performed a genome-wide CRISPRi screen in human cells to identify genes that influence the import of proteins targeted to peroxisomes. In addition to known *PEX* genes, we found that knockdown of the E3 ligase RNF146 reduces import of PTS1-tagged proteins into the peroxisome. RNF146 (Ring Finger Protein 146), also known as Iduna, is a RING-domain E3 ubiquitin ligase that recognizes and ubiquitinates proteins modified by poly(ADP-ribosyl)ation (PARsylation) [Zhang et al. 2011, DaRosa et al. 2015]. RNF146 interacts directly with poly(ADP-ribose) polymerases, such as TNKS and TNKS2 [Da Rosa et al. 2015] and PARP1 and PARP2 [Gero et al 2014, Kang et al 2011]. Together, the poly(ADP-ribose) polymerases and RNF146 specifically regulate the stability of numerous substrates which are first PARsylated and subsequently polyubiquitinated by RNF146, triggering proteasomal degradation. We found that RNF146-mediated loss of peroxisomes was dependent on the accumulation of the poly(ADP-ribose) polymerase tankyrase, specifically by impairing import into peroxisomes through a mechanism dependent on tankyrase’s activity as a poly(ADP-ribose) polymerase. We thus propose a model in which tankyrase binds and PARsylates PEX14 and neighboring proteins, inhibiting the import of PTS1-tagged proteins.

RNF146 and tankyrase are better known as co-regulators of protein stability: tankyrase binds and PARsylates substrates with a tankyrase-binding motif, which then triggers poly-ubiquitination by RNF146 [DaRosa et al 2015]. Known TNKS/RNF146 substrates include AXIN1, BLZF1, 3BP2, and CASC3 [Nie et al 2020, Levaot et al 2011]. Surprisingly, we found that in a variety of cell lines, a loss of *PEX* genes altered the stability of RNF146/TNKS substrates and could therefore alter the output of downstream signaling pathways, including the Wnt/beta-catenin pathway. These observations suggest that not only is peroxisome abundance and function integrally intertwined with cell signaling pathways, but also that peroxisomes themselves regulate cellular responses to external stimuli.

## Results

### Sequestration of ZeoR in peroxisomes links peroxisome import to viability

Past screens for peroxisomal genes in mammalian cells have relied on peroxisome-localized enzymatic activity [Zoeller and Raetz 1986, Tsukamoto et al 1990; Morand et al. 1990] and fluorescence microscopy of PTS1-tagged fluorescent proteins [Ito et al 2000], since mammalian cells in tissue culture conditions do not require peroxisomes for growth. To facilitate a CRISPRi screening approach for regulators of peroxisome function, we engineered a cell line, which we term Pex-ZeoR, in which the efficiency of peroxisome import is linked to cell viability by fusing the fluorescent marker mVenus and a peroxisomal targeting signal (PTS1) to the gene encoding resistance to Zeocin, a 1400 Dalton molecule in the bleomycin family that induces DNA double strand breaks and causes cell death [Murray et al 2014; Drocourt et al 1990]. With this fusion construct, mVenus-ZeoR-PTS1, cells with functional peroxisomes should sequester the Zeocin resistance protein (ZeoR), thereby preventing them from neutralizing Zeocin, which is too large to passively diffuse through peroxisome membranes [Antonenkov and Hiltunen 2006]. By contrast, cells with reduced peroxisome import should accumulate mVenus-ZeoR-PTS1 in the cytoplasm where it can neutralize Zeocin, conferring a selective advantage in the presence of Zeocin (**Fig. 1A**). To affirm our strategy, we transduced HCT116 CRISPRi (dCas9-KRAB) cells [Liang et al 2018; Gilbert et al 2014] to recombinantly express mVenus-ZeoR-PTS1. As predicted, cells expressing a non-targeting control (NTC) sgRNA had fluorescent mVenus foci, while cells expressing a *PEX1* targeting sgRNA exhibited diffuse cytosolic mVenus signal (**Fig. 1B**), consistent with mVenus-ZeoR-PTS1 targeting to the peroxisome. We then assessed cell growth of the HCT116 CRISPRi Pex-ZeoR cell line over a range of Zeocin concentrations, finding a clear growth advantage for cells with sgRNAs targeting *PEX1* or *PEX6* versus NTC at high concentrations of Zeocin (**Fig S1A**). To identify optimal selection conditions for the genome-wide screen, we performed a competition assay by co-culturing either PEX1 and NTC or PEX6 and NTC CRISPRi Pex-ZeoR cells at varying dosages of Zeocin, and monitoring the abundance of each cell population by flow cytometry. While *PEX1* and *PEX6* knockdown cells were outcompeted by NTC cells in conditions without Zeocin, they displayed a marked competitive advantage even at low concentrations of Zeocin (**Fig. 1C, Fig S1B**). Together, these validation experiments suggest that peroxisomal sequestration of ZeoR allows for the selection of cells harboring sgRNAs that target peroxisomal genes.

**Figure 1.**
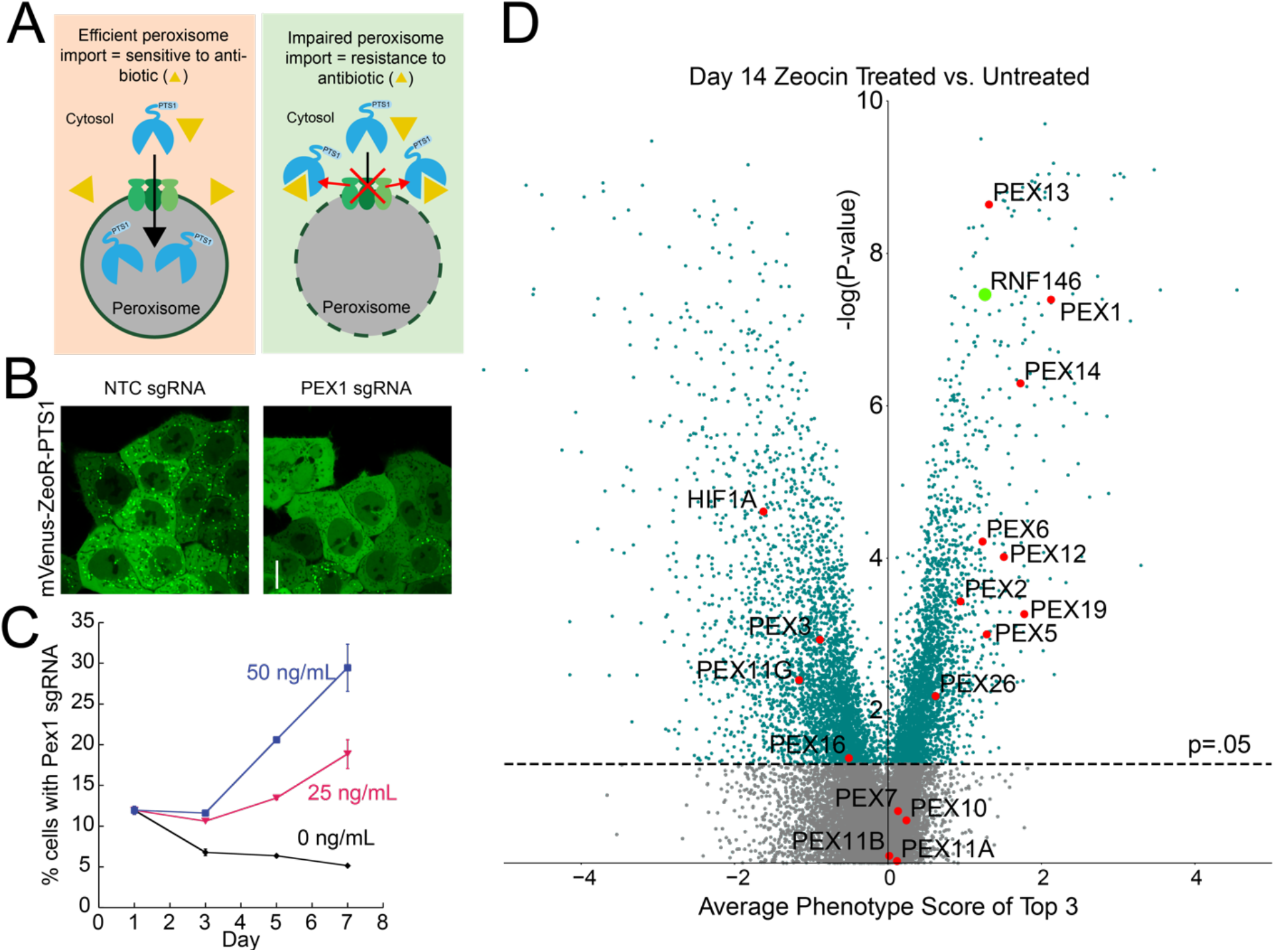
A genome-wide screen uncovers genes that regulate peroxisome biology. **(A)** Design of the Pex-ZeoR cell line, which sequesters the Zeocin resistance protein in the peroxisome matrix. Loss of PEX genes causes cytosolic Zeocin resistance. **(B)** Representative fluorescence microscopy images of live HCT116 mVenus-ZeoR-PTS1 cells expressing either NTC or PEX1 sgRNAs. Fusion construct forms puncta in WT but not aperoxisomal (PEX1 knockdown) cells. Fluorescent microscopy data are representative of n=49 images from m=2 biological replicates. Scale bar: 10μm **(C)** Quantification of flow cytometry data of BFP- (NTC) and BFP+ (PEX1) cells grown in co-culture competition assay over t=11 days in the presence of 0, 25, or 50 ng/uL of Zeocin. Timepoints are taken every t=2 days. Data shown as the mean ± SD of n=3 biological replicates. **(D)** Volcano plot of NGS data from genome-wide screen with significance (-log base 10 of p-value, y-axis) and phenotype score (normalized fold change of cDNA guide count, x-axis) of guides targeting specific genes for cell cultures either untreated (DMSO mock treated) or treated (50ng/uL Zeocin treated) for 14 days. Teal data points are below a cut-off of p=0.05. Red data points represent known PEX genes and HIF1A. The green datum point represent RNF146. Data displayed was calculated from m=3 guides per gene and n=2 biological replicates.

### A genome-wide CRISPRi screen in Pex-ZeoR cells enriches known *PEX* genes

Emboldened, we executed a genome-wide screen with the Pex-ZeoR cell line to identify novel genes that affect peroxisomal homeostasis. Infection with a genome-wide CRISPRi library was followed by chronic treatment with or without Zeocin, combined with regular passaging of cells over 35 days, with samples collected every 7 days for terminal Illumina sequencing preparation (**Fig. S1C**). We found 1,719 genes that were significantly different (*p*<0.05) between the treated and untreated conditions at the day 14 timepoint (**Fig. 1D**). Day 14 serves as the optimal comparison timepoint because of clear enrichment of the majority of known *PEX* genes while maintaining sufficient library diversity and replicate quality (**Fig S1D, S1E**).

We observed enrichment of guides targeting known *PEX* genes that facilitate PTS1 import (*PEX5, PEX13, PEX14, PEX2/PEX12, PEX1/PEX6, PEX26*) and peroxisome membrane protein targeting (*PEX19*) affirming the efficacy of our strategy (**Fig. 1D**). Guides targeting *PEX7* and alpha and beta variants of *PEX11* were not strongly enriched, consistent with roles in recognition of the alternative PTS2 targeting signal (*PEX7*) [Braverman et al 1997], and peroxisomal membrane elongation (*PEX11*) [Koch et al 2010]. Perplexingly, guides targeting one component of the peroxisome RING finger complex, PEX10, were not enriched compared to the other constituents, PEX2 and PEX12, and guides targeting other peroxisome membrane biogenesis factors PEX3 and PEX16, were depleted in the screen (**Fig 1D, S1E**). Of the known factors regulating peroxisome specific autophagy, such as NBR1, MARCH5, SQSTM1, HIF1A, and NIX [Kim et al 2008, Deosaran et al 2013, Zheng et al 2022, Wilhelm et al 2022], we found that only guides targeting *HIF1A*, the loss of which stabilizes peroxisomes [Wilhelm et al 2022], were strongly depleted in our screen. Although most peroxisome-homeostasis related genes behaved according to our predictions, a handful did not align with our a priori prognosis. Our results suggest the possibility that not all of the aforementioned genes are simple or monotonic in their effect on peroxisome import or autophagy, representing potential new mechanisms for further investigation.

### Guides targeting *RNF146, INTS8, KCNN4* reduce peroxisomal foci intensity

We anticipated that sgRNAs that improve resistance to Zeocin independent of the peroxisomal localization of ZeoR should also be significantly enriched in our dataset. Thus, to narrow the candidate list to genes relevant to peroxisomal localization of ZeoR, we filtered our screen results to exclude factors that modulated resistance to a related DNA damaging agent, bleomycin [Olivieri et al 2020] (Z-score range [-0.5,0.5]). GO analysis of the remaining genes with a fold change greater than 2 and a Mann-Whitney p<0.05 revealed a 100-fold enrichment of GO terms related to protein import into the peroxisome, and a greater than 50-fold enrichment related to RNA cleavage involved in mRNA processing. We note that several *PEX* genes (*PEX1, PEX6, PEX12*) modulate bleomycin resistance, possibly because there is a direct link between DNA repair and peroxisome biology through localization of the DNA repair kinase ATM to peroxisome membranes [Zhang et al 2015].

We then used fluorescence microscopy of mVenus-PTS1 in the Pex-ZeoR cell line to assess how knockdown of candidate genes altered peroxisome abundance. For each candidate gene, we produced two unique constitutive knockdown cell lines per gene and quantified mVenus-PTS1 foci number, foci and cell area, and foci and cytoplasm fluorescence intensity using CellProfiler [Stirling et al. 2021]. To estimate the efficiency of peroxisome import while accounting for different mVenus-PTS1 expression levels, we calculated the ratio of the intensity of mVenus-PTS1 in peroxisome foci to the intensity of mVenus-PTS1 in the cytoplasm (**Fig 2A, S2A, S2B**). We found that several of the guides enriched by Zeocin selection decreased the ratio of peroxisomal to cytosolic mVenus-PTS1 intensity, including those targeting the E3 ligase RNF146, Integrator complex subunit INTS8, and calcium-activated potassium channel KCNN4.

**Figure 2.**
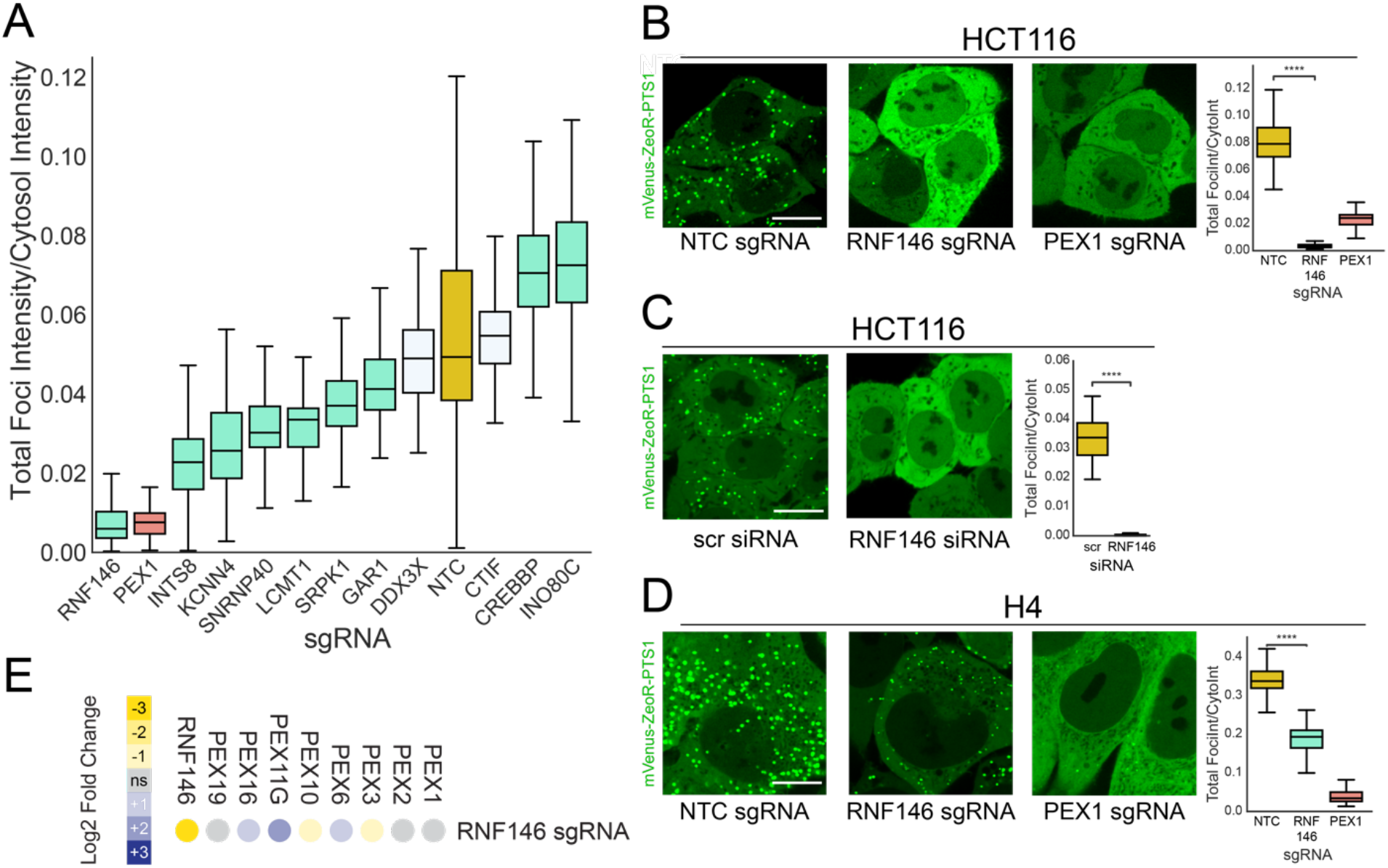
Peroxisome abundance is regulated locally by RNF146. **(A)** CellProfiler quantification of the ratio of mVenus-PTS1 intensity in foci and in the cytoplasm in fluorescence microscopy images acquired of live HCT116 Pex-ZeoR cells expressing sgRNAs targeting various genes. Data per gene constitutes m=2 unique sgRNAs with n=49 images per gene. Non-targeting control sgRNA shown in yellow, PEX1 sgRNA shown in pink, sgRNAs significantly different from NTC (p<0.0001, independent t-test) are in blue, sgRNA with p>0.05 are in white. **(B)** Representative fluorescence microscopy images of mVenus expression in HCT116 Pex-ZeoR cells harboring sgRNAs for NTC, PEX1, or RNF146 and quantification of the ratio of mVenus-PTS1 foci intensity to mVenus-PTS1 cytosolic intensity. Data is representative of m= 49 images. n=2 biological replicates. Scale bars: 10 μm. **(C)** Representative fluorescence microscopy images of mVenus expression in HCT116 Pex-ZeoR cells treated with either scrambled (scr) siRNA or RNF146 siRNA, and quantification of the ratio of total mVenus-PTS1 foci intensity to mVenus-PTS1 cytosolic intensity. Data is representative of m= 49 images n=2 biological replicates. Scale bar: 10 μm. **(D)** Representative fluorescence microscopy images of mVenus expression in H4 Pex-ZeoR cells harboring sgRNAs for NTC, PEX1, or RNF146 and quantification of the ratio of mVenus-PTS1 foci intensity to mVenus-PTS1 cytosolic intensity. Data is representative of m= 49 images. n=2 biological replicates. Scale bars: 10μm. Asterisks denote p <0.01, ***p <0.001, ****p <0.0001, whereas ns denotes not significant, calculated by independent t-test. **(E)** Heatmap of RNA-seq data displaying significant (p<0.05) fold change of *PEX* gene transcription in RNF146 knockdown cells versus NTC controls. Data is representative of n=3 biological replicates.

### RNF146 regulates peroxisome foci intensity in multiple cell lines

Given the magnitude of the impact of the RNF146 knockdown on mVenus-PTS1 foci (**Fig. 2A, 2B**), we chose to focus our efforts on characterizing the effects of RNF146 on peroxisome homeostasis. We first ruled out possible off-target effects of the RNF146 sgRNA by treating our reporter cell line with RNF146 siRNA, which recapitulated the loss of mVenus foci signal within 24 hours of siRNA treatment (**Fig. 2C**). To determine if the peroxisomal effect of RNF146 knockdown was specific to the HCT116 cell line, we created a secondary cell line, the H4 astrocytoma cancer cell line, harboring the same CRISPRi machinery and our Pex-ZeoR reporter. We observed significant depletion of mVenus-PTS1 foci intensity in both the HCT116 and H4 *RNF146* and *PEX* knockdown cell lines (**Fig. 2B, 2D**). The significant depletion of PTS1 foci in two independent cell lines suggests that RNF146 has a bona fide role in regulating peroxisome homeostasis in human cells.

To determine if *RNF146* KD impacted peroxisome biogenesis through an effect on *PEX* gene expression, we gathered RNA-seq data of *RNF146* KD HCT116 cell mRNA transcripts versus NTC cells. We found that knockdown of *RNF146*, which was confirmed in the data set, mildly repressed transcription of *PEX3* and *PEX10*. Given that neither *PEX3* nor *PEX10* had positive phenotype scores in the CRISPRi screen, we found it unlikely that the RNF146 phenotype can be completely explained by these transcriptional changes, thereby indicating a post-transcriptional role for RNF146 in regard to peroxisomal homeostasis (**Fig 2E**).

### RNF146-mediated loss of mVenus-PTS1 foci depends on TNKS, but not autophagy

RNF146 is known to collaborate with poly(ADP-ribose) polymerases to ubiquitinate PARsylated proteins and target them for degradation. Loss of RNF146 is therefore expected to stabilize PARsylated substrates, which could act to either inhibit peroxisome biogenesis or increase peroxisome-specific autophagy. We therefore tested if the observed loss of mVenus-PTS1 foci in response to RNF146 knockdown depended on changes in the RNF146 partner, TNKS. We first assessed TNKS levels in an RNF146 knockdown, and found that knockdown of RNF146 expression in the HCT116 Pex-ZeoR cell line caused a marked increase in TNKS protein levels (**Fig 3A**). To test if RNF146’s effect on peroxisomes depended on increased TNKS levels, we performed a dual knockdown assay of *RNF146* and *TNKS* in our reporter cell line. We found that siRNA knockdown of *TNKS* in RNF146 CRISPRi cells rescued the import of mVenus-PTS1 (**Fig 3B**) indicating that RNF146’s effect on peroxisomes depended on TNKS. These results are consistent with previous reports that TNKS is significantly stabilized in cells lacking RNF146 [Nie et al 2020]. Although it was previously shown that TNKS mediates peroxisome-specific autophagy [Li et al 2017], we found that siRNA inhibition of ATG7 did not prevent the accumulation of TNKS nor the loss of mVenus-PTS1 foci intensity in RNF146 knockdown cells **(Fig 3C, 3D**). This lack of dependence on autophagy was further corroborated in multiple cell lines by the treatment of RNF146 knockdown cells with autophagy inhibitors bafilomycin or hydroxychloroquine, which, despite preventing LC3BII turnover, did not substantially rescue peroxisome foci number or intensity relative to control cells **(Fig S3A-E)**. These observations suggest that while the effect of RNF146 knockdown on peroxisomes depends on TNKS, it does not depend on peroxisome-specific autophagy.

**Figure 3.**
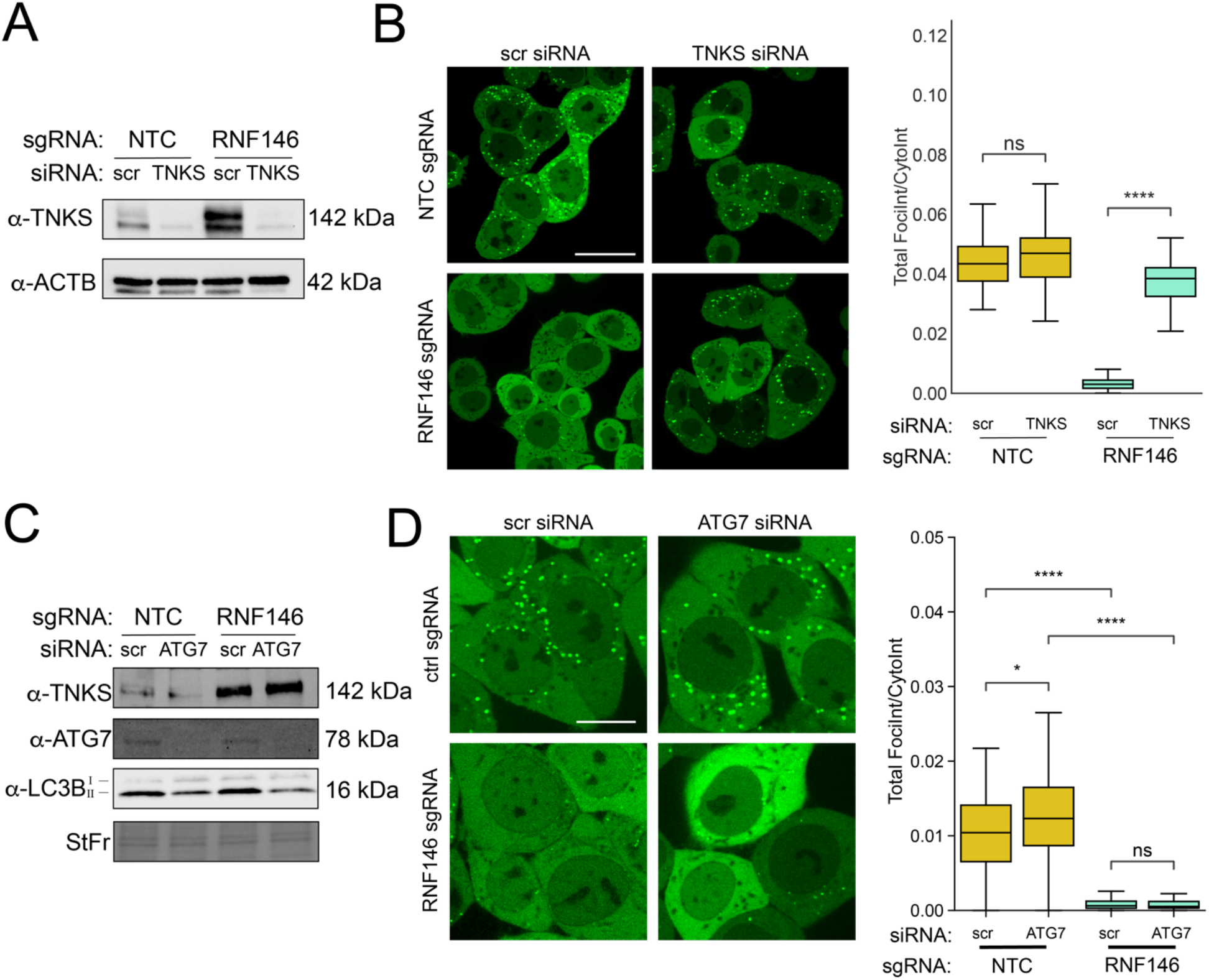
RNF146’s effect on peroxisomes is mediated by tankyrase, but not autophagy. **(A)** Immunoblots of HCT116 Pex-ZeoR cells with antibodies against TNKS and ACTB (loading control). **(B)** Left panel: Representative mVenus-PTS1 fluorescence microscopy images of either non-targeting control (NTC) or RNF146 sgRNA cells treated with either scrambled (scr) siRNA or TNKS siRNA. Right panel: Quantification of mVenus-PTS1 microscopy images in left panel for mVenus-PTS1 foci intensity versus total cytosol intensity in HCT116 Pex-ZeoR cells. Data is representative of 49 images per condition and 2 biological replicates. Scale bars: 10 μm. **(C)** Immunoblot for TNKS, ATG7, and LC3B in lysate from scrambled or ATG7 siRNA treated HCT116 Pex-ZeoR cells with sgRNAs for either NTC or RNF146. **(D)** Left panel: Fluorescence microscopy data of scrambled or ATG7 siRNA treated HCT116 Pex-ZeoR cells with sgRNAs for either non-targeting control (NTC) or RNF146. m=32 images. n=2 biological replicates. Right panel: Quantification of mVenus-PTS1 microscopy images for mVenus foci intensity versus cytosol intensity in HCT116 Pex-ZeoR cells. Scale bars: 10μm. All blots are representative of n=3 biological replicates. Asterisks denote p-values *p <0.05, **p <0.01, ***p <0.001, ****p <0.0001, whereas ns denotes not significant, calculated by independent t-test.

### Loss of RNF146 specifically inhibits import into peroxisomes

Since the loss of RNF146 did not appear to induce peroxisome-specific autophagy, we evaluated whether the loss of RNF146 could specifically impair peroxisome biogenesis at the stage of protein import into peroxisomes. We performed immunofluorescence microscopy on the HCT116 and H4 CRISPRi Pex-ZeoR cell lines harboring sgRNAs for *NTC, RNF146, PEX5*, and *PEX19*, where PEX5 and PEX19 are the receptors for PTS1-tagged matrix protein import and peroxisomal membrane protein insertion, respectively **(Fig. 4A, S4A**). We found that knockdown of RNF146 in both HCT116 and H4 cells resembled a PEX5 knockdown, in which a peroxisome membrane protein PMP70 remains present and punctate **(Fig 4A, 4B, S4A, S4B**), but matrix proteins, both mVenus-PTS1 and catalase, no longer form foci **(Fig 4A, 4C, Fig. S4A, S4C**) or co-localize with PMP70 **(Fig S4D**). These observations suggest that loss of RNF146 specifically inhibits import of PEX5 client proteins into the peroxisome.

**Figure 4.**
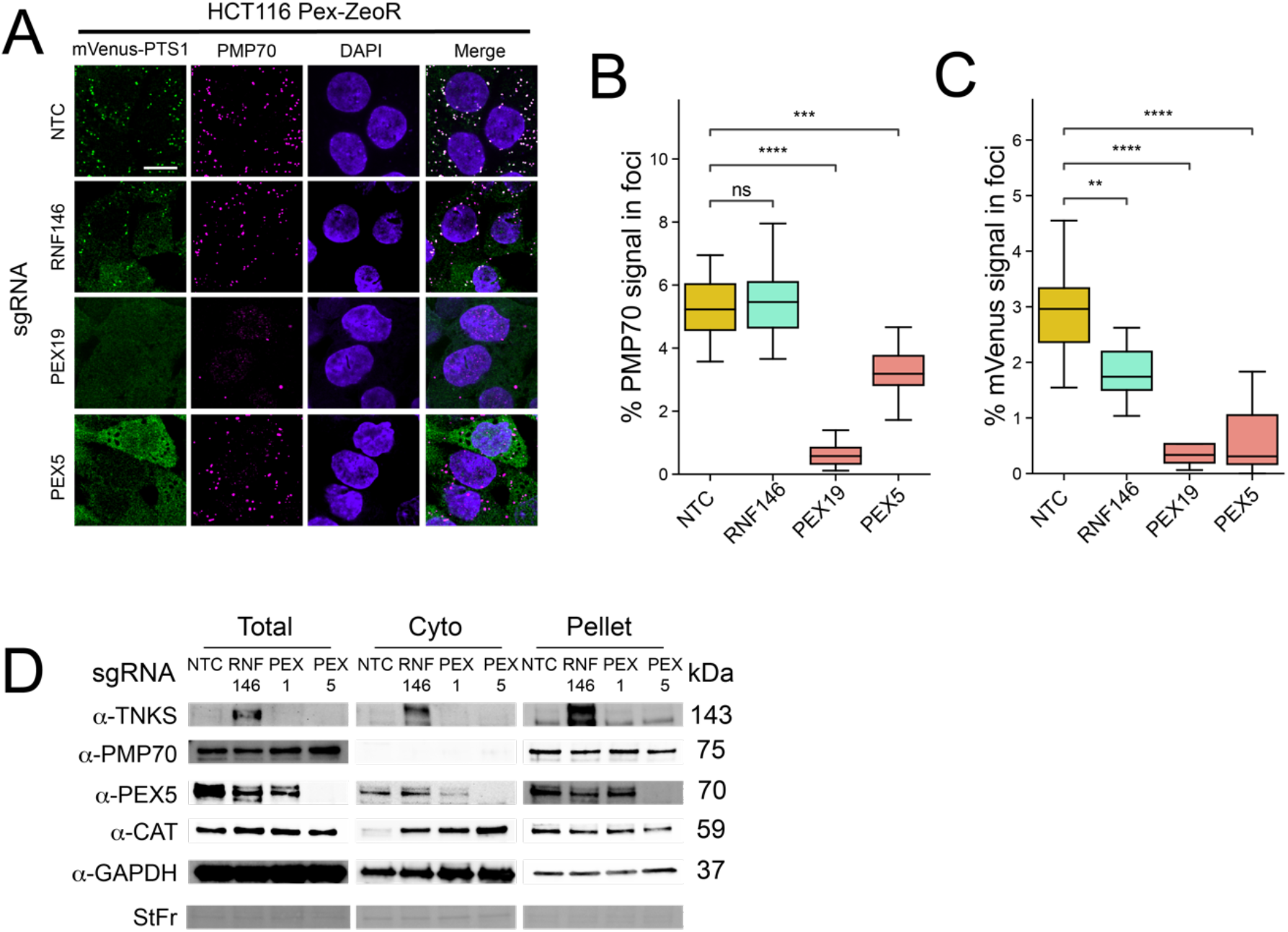
Loss of RNF146 impairs peroxisome protein import. **(A)** Representative immunofluorescence microscopy images of NTC, RNF146, PEX19, and PEX5 sgRNA expressing HCT116 Pex-ZeoR cells. mVenus-PTS1 in green, DAPI in blue, PMP70 in magenta. **(B, C)** Quantification of immunofluorescence microscopy images for percentage foci area of PMP70 (B) and mVenus-PTS1 (C) versus cytosolic area. m=25 images. n=2 biological replicates. **(D)** Immunoblot of HCT116 Pex-ZeoR with sgRNAs targeting NTC, RNF146, PEX1, PEX5. Fractions represent total lysate, 20,000xg supernatant, and 20,000xg pellet. All blots are representative of n=3 biological replicates. Asterisks denote p-values *p <0.05, **p <0.01, ***p <0.001, ****p <0.0001, whereas ns denotes not significant, calculated by independent t-test.

Efficient peroxisomal matrix protein import relies on PEX5 binding to the PTS1-tagged protein, PEX5 docking to PEX13/PEX14 at the peroxisome, and extraction of ubiquitinated PEX5 from the peroxisome membrane by the PEX1/PEX6/PEX26 motor complex for continued rounds of import. PEX5 is therefore typically distributed between both cytoplasmic and membrane fractions, with an increased proportion at the peroxisome membrane in mutants of the ubiquitination and extraction machinery [Platta et al 2005]. To determine if RNF146 knockdown alters the localization of PEX5, we probed for PEX5 and catalase in soluble and membrane fractions after fractionation. As expected, we observed that PEX5 distributes between both membrane and soluble fractions in wild type cells, and that more PEX5 is trapped in the membranes due to impaired extraction of PEX5 in PEX1 knockdown cells. Interestingly, a larger proportion of PEX5 was soluble in RNF146 knockdown cells compared to controls cells **(Fig. 4D)**. This suggests that the impairment of import of peroxisomes may be due to reduced recruitment of PEX5 and PTS1-cargo to the peroxisome membrane. Additionally, we observed that the soluble proportion of catalase, a PEX5 client protein without a canonical PTS1 tag, increased in RNF146, PEX5, and PEX1 knockdown cells, confirming that RNF146 knockdown also impedes import of endogenous matrix proteins **(Fig. 4D)**.

### PARP activity of TNKS impedes import into peroxisomes

TNKS contains N-terminal ankyrin repeats that bind substrates with a tankyrase binding motif, a SAM domain that mediates oligomerization, and a C-terminal poly(ADP-ribose) polymerase domain [Guettler et al 2011]. There are predicted, conserved tankyrase binding motifs in PEX14, PEX5, PEX19, and PEX11G [Guettler et al 2011]. We found that TNKS co-immunoprecipitated both FLAG-PEX14 and PEX5 upon RNF146 knockdown **(Fig. 5A)**. Additionally, FLAG-PEX14 co-immunoprecipitated TNKS and PEX5 in NTC and RNF146 knockdown cells **(Fig. 5B)**. These results suggest TNKS associates with the peroxisome membrane and peroxisome import machinery upon RNF146 knockdown.

**Figure 5.**
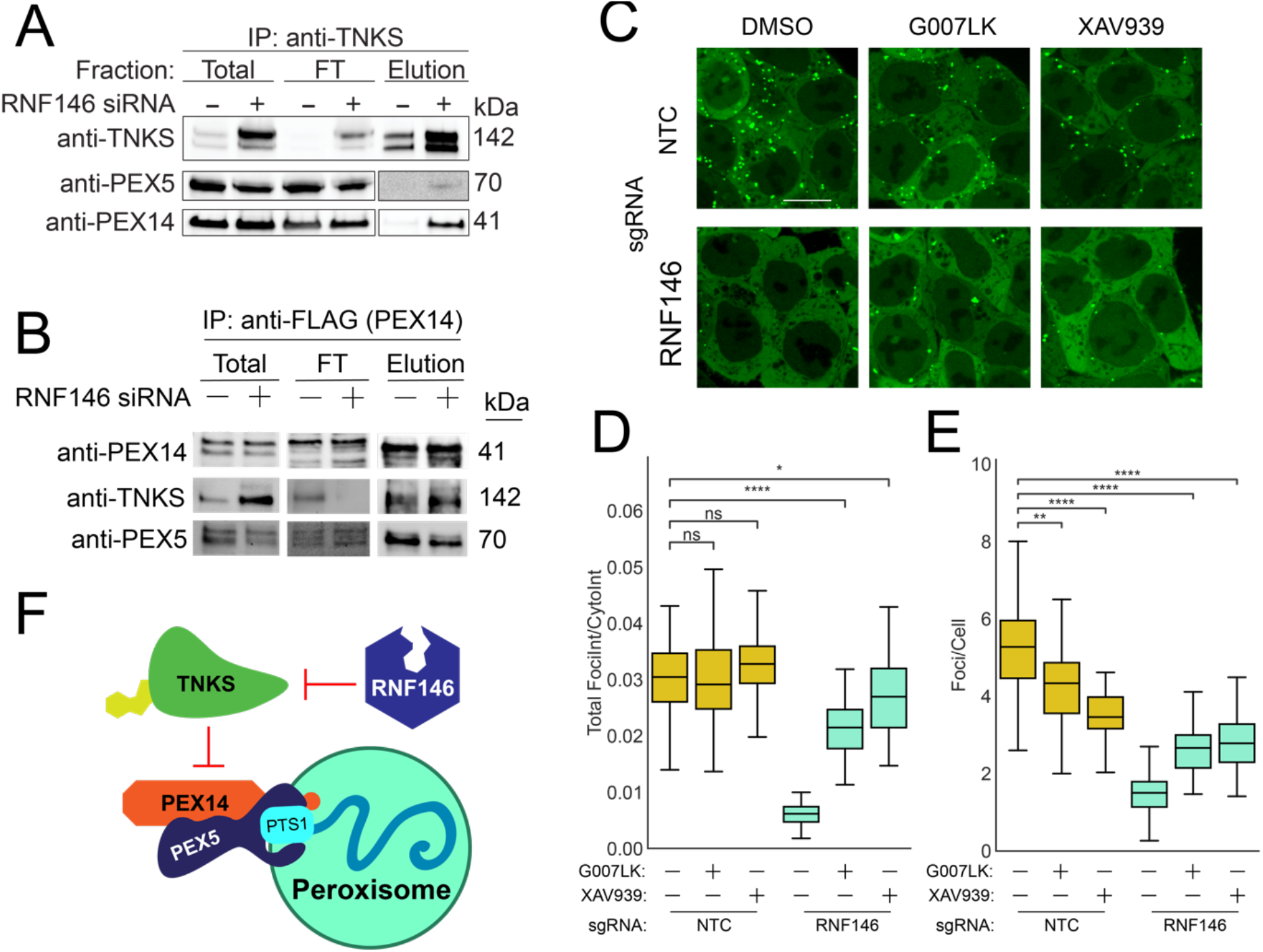
TNKS PARP activity impairs peroxisome protein import. **(A)** Immunoblots of anti-TNKS immunoprecipitation fractions from HCT116 Pex-ZeoR cells expressing PEX14 sgRNAs with constitutive re-expression for FLAG-PEX14 treated with either NTC or RNF146 siRNA (10 nM) for 24hrs, detecting TNKS, PEX5, and PEX14. **(B)** Immunoblots of anti-FLAG immunoprecipitation fractions from HCT116 Pex-ZeoR cells expressing PEX14 sgRNAs with constitutive re-expression of FLAG-PEX14, treated with either NTC or RNF146 siRNA (10 nM) for 24hrs, detecting TNKS, PEX5, and PEX14. **(C)** Representative live-cell fluorescence microscopy images of NTC and RNF146 sgRNA expressing HCT116 Pex-ZeoR cells treated with DMSO (mock), 500 nM G007LK, or 10 μM XAV939 for 24 hrs. mVenus-PTS1 in green. Scale bar: 10 μm. **(D)** Quantification of fluorescence microscopy images for the ratio of mVenus- PTS1 foci intensity to mVenus-PTS1 cytosolic intensity (left) and the number of foci per cell (right). m=32 images and n=2 biological replicates. All blots are representative of n=3 biological replicates. Asterisks denote p-values *p <0.05, **p <0.01, ***p <0.001, ****p <0.0001, whereas ns denotes not significant, calculated by independent t-test. (**E**) Proposed model: loss of RNF146 increases active TNKS, which binds PEX14 and PARsylates proteins at the peroxisome membrane impairing peroxisome import.

To test if RNF146’s effect on peroxisomes depended on the PARP activity of TNKS, we tested if the TNKS1/2 inhibitors G007LK and XAV939 restored peroxisome foci in RNF146 knockdown cells **(Fig 5C, 5D)**. We found that TNKS inhibitors partially restored import of mVenus-PTS1 into foci in RNF146 knockdown cells as judged by the ratio of foci to cytosolic intensity of mVenus-PTS1 (**Fig 5D**), but did not fully recover peroxisome number **(Fig 5E)**. This observation suggests that tankyrase’s PARsylation activity is important for RNF146’s effect on peroxisomes. We therefore propose a model in which high levels of active TNKS, induced by loss of RNF146, binds PEX14 and PARsylates proteins at the peroxisome membrane, which inhibits PEX5-mediated protein import into peroxisomes (**Fig. 5E)**.

### PEX proteins alter RNF146/TNKS activity towards other substrates

This model suggests that TNKS binds peroxisome membrane protein PEX14 and can localize to the peroxisome. Other better-known substrates of TNKS, such as BLZF1, which localizes to the Golgi [Yue et al 2021], and AXIN1, which localizes to centrosomes [Lach et al 2022], have defined locations elsewhere in the cell. We thus wondered if peroxisomal recruitment of TNKS could regulate access to other substrates. To test if the presence of peroxisome membranes and membrane proteins alters TNKS substrate selection, we evaluated the stability of the TNKS/RNF146 substrates AXIN1, CASC3, and BLZF1 in cells with knockdown of the peroxisomal membrane protein *PEX14*, the peroxisomal membrane protein chaperone *PEX19*, or a non-targeting control. We found that AXIN1 and CASC3 levels were significantly depleted in *PEX19* knockdown HCT116 cells, and BLZF1 levels were depleted in both *PEX19* and *PEX14* knockdown HCT116 cells (**Fig. 6A**). Furthermore, *PEX14* and *PEX19* knockdowns also depleted AXIN1 levels in HEK293T, iPSC AICS-0090-391, and H4 CRISPRi cells (**Fig. 6B, 6C, Fig. S5A**), illustrating that this phenomenon is not specific to HCT116 cells. To confirm that the effect of *PEX19* knockdown arises from loss of PEX19, we re-expressed PEX19 using a lentiviral vector to complement the knockdown of endogenous *PEX19*, and observed a rescue of AXIN1 stability (**Fig. 6D**). Additionally, suppression of either *RNF146* or *TNKS* mRNA transcripts via siRNA, as well as XAV939 mediated catalytic inhibition of TNKS, restored AXIN1 stability in *PEX19* knockdown cells, demonstrating that loss of PEX19 activates RNF146/TNKS-mediated destabilization of AXIN1 (**Fig. 6D**). These observations suggest that functional peroxisomes repress TNKS activity towards some substrates, including AXIN1, BLZF1, and CASC3.

**Figure 6.**
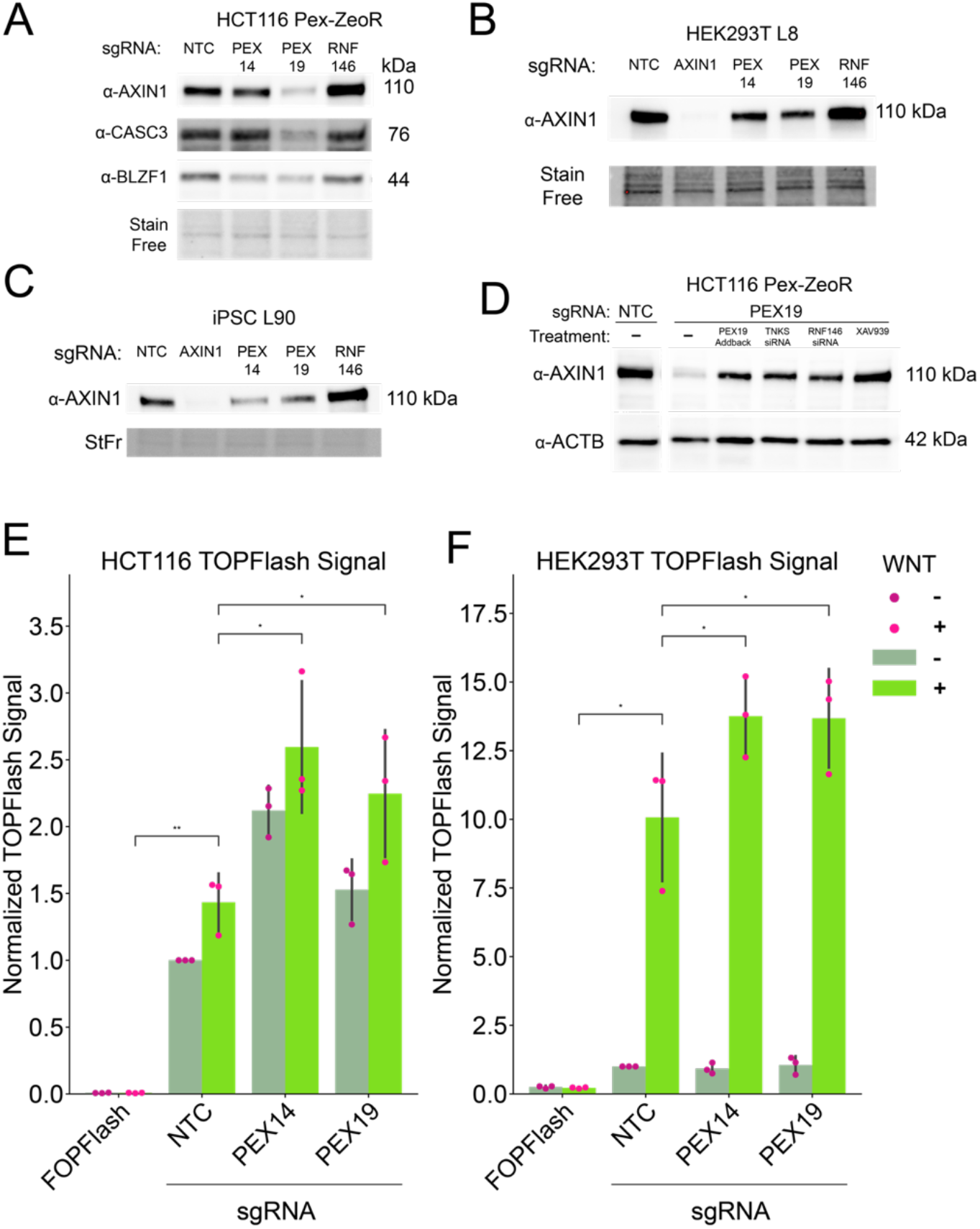
Peroxisome abundance influences of RNF146/TNKS substrate selection. **(A-C)** Immunoblots measuring the abundance of AXIN1, CASC3, and BLZF in **(A)** HCT116 (n=3), **(B)** HEK293 (n=3), and **(C)** iPSC AICS-0090-391 (n=3) CRISPRi cells with indicated sgRNAs. **(D)** Western blot measuring abundance of AXIN1 and ACTB (loading control) in HCT116 cells with indicated sgRNAs, PEX19 knockdown cells are paired with treatments for PEX19 reexpression, TNKS siRNA (10 nM), RNF146 siRNA 10 nM), or XAV939 (10 μM). Blots shown are representative of n=1 blots. **(E-F)** TOPFlash Dual Luciferase assays measuring the induction of Wnt signaling to downstream beta-catenin transcription in PEX knockdown HCT116 cells (E) and HEK293T (F) harboring the indicated sgRNAs and treated with or without 315ng/mL Wnt3a for 24 hrs (data shown is 48 hours post-transfection with TOPFlash constructs). Luciferase activity is measured versus a renilla transfection control and data is normalized to untreated NTC samples. FOPFlash negative control performed in NTC sgRNA cells. Data is representative of n=3 biological samples. Asterisks denote p-values *p <0.05, **p <0.01, ***p <0.001, ****p <0.0001, whereas ns denotes not significant, calculated by paired t-test.

### Increased Wnt/beta-catenin signaling in *PEX* knockdown cells

AXIN1 is the limiting component for the formation of the beta-catenin destruction complex which induces the phosphorylation and subsequent degradation of the beta-catenin transcription factor. In canonical Wnt signaling, Wnt ligand binding to the Frizzled receptor dissociates the beta-catenin destruction complex, allowing beta-catenin to accumulate, enter the nucleus, and induce transcription of Wnt-responsive genes. The stabilization of AXIN1, such as by TNKS inhibitors, inhibits Wnt signaling by increasing levels of the destruction complex [Huang et al 2009]. Since AXIN1 was severely destabilized in *PEX19* knockdown HCT116 cells and partially destabilized in *PEX14* and *PEX19* knockdown HEK293T, H4, and iPSC AICS-0090-391 cells, we tested if the knockdown of *PEX* genes can therefore influence the Wnt signaling pathway using the TOPFlash reporter for beta-catenin transcriptional activity. We found that HCT116 cells had a greater transcriptional response to Wnt ligand in *PEX14* and *PEX19* knockdown cells (**Fig. 6E**), as well as increased basal activity. Since HCT116 cells are derived from a colorectal carcinoma heterozygous for a dominant mutation in beta-catenin that causes constitutively active beta-catenin-TCF regulated transcription [Morin et al 1997], we also tested the effect of the PEX knockdowns on the TOPFlash reporter in HEK293T cells. Both *PEX14* and *PEX19* knockdown HEK293Ts exhibited a partial loss of AXIN1 levels (**Fig. 6B**), and consistently, also exhibited a greater response to Wnt ligand, though basal levels were not perturbed (**Fig. 6F**). Our observations show that knockdown of *PEX14* and *PEX19* increases Wnt signaling consistent with the decreased levels of the core subunit of the beta-catenin destruction complex, AXIN1.

## Discussion

Here we describe an approach to link cell viability to peroxisome import efficiency by sequestering the Zeocin resistance protein in the peroxisome. We use this approach to screen for novel genes regulating peroxisome import efficiency. In addition to known PEX genes, we found that the E3 ligase RNF146 regulates peroxisome import through its control of the levels of the poly(ADP-ribose) polymerase tankyrase. High levels of tankyrase, which can bind PEX14 and possibly other PEX proteins, specifically inhibits import into peroxisomes. In our cell lines, inhibition of import depends on tankyrase’s poly(ADP-ribose) polymerase activity. We note that Li and colleagues showed that increased levels of tankyrase due to treatment with tankyrase inhibitors such as XAV939 could induce peroxisome-specific autophagy in HEK-293T cells; however, this autophagy does not mediate the loss of mVenus-PTS1 foci in response to RNF146 knockdown in H4 or HCT116 cells. Instead, we find that tankyrase’s polymerase activity is required for the observed inhibition of peroxisome import. We therefore propose a model in which loss of RNF146 stabilizes active tankyrase, which PARsylates proteins at the peroxisome membrane and impairs their function in matrix protein import into the peroxisome.

This model suggests that any mechanism that inactivates RNF146 will inhibit import into peroxisomes. In mice, RNF146 transcription is repressed during RANKL-mediated osteoclastogenesis through an NF-κB binding site [Matsumoto et al 2017], suggesting that peroxisome import may be coordinated with cell type specification through RNF146 and tankyrase. RNF146 activity is also regulated by sumoylation [Li et al 2023], localization to the nucleus [Gero et al 2014; Sheng et al 2018], and direct interaction with other poly(ADP-ribose) polymerases such as PARP-1 [Gero et al 2014]. It is therefore possible that temporary localization of RNF146 to the nucleus in response to DNA damage could impede peroxisome import, perhaps to increase concentrations of cytosolic catalase to reduce oxidative stress. The effect of this regulation on tankyrase activity and peroxisome import, and the consequences for RNF146’s protective role during DNA damage [Kang et al 2011], oxidative stress [Xu et al 2013], and PARsylation induced cell death [Andrabi et al 2011] warrants further investigation.

A second implication of our results is that RNF146/TNKS together may regulate the stability of substrates at the peroxisome membrane, such as PEX14 itself, or neighboring proteins. Proteomic studies show that the loss of TNKS significantly stabilizes PEX14 and SLC27A2, a peroxisomal transporter for long chain fatty acids [Bhardwaj et al 2017]. We did not observe a change in peroxisome protein import or peroxisome number in response to TNKS knockdown (**Fig 2B**). However, it is possible that this may be due to the relatively low levels of expression of endogenous TNKS in the HCT116 cell line, and in cells with high levels of TNKS, such as the brain, adipose tissue, and endocrine pancreas [Yeh et al 2009], it is possible that a knockdown of TNKS could improve peroxisome import and abundance. Indeed, studies of TNKS deficient mice show that they have increased fatty acid oxidation, which is consistent with improved peroxisomal function [Yeh et al 2009]. It is also possible that RNF146/TNKS activity at the peroxisome membrane regulates signaling from the peroxisome membrane. For example, RNF146/TNKS coordinate the degradation of the antiviral protein MAVS [Xu et al 2022], which has been shown to localize to both the peroxisome and mitochondria and initiate disparate signaling pathways upon viral infection [Dixit et al 2010].

An intriguing corollary of RNF146/TNKS localization to the peroxisome membrane is the impact of this localization on its access to other substrates. We found that the knockdown of different PEX proteins, particularly PEX14, which binds TNKS, and PEX19, which is generally required for peroxisome membrane protein stability, decreases the stability of RNF146/TNKS substrates that are not thought to be at the peroxisome. We propose that localization to the peroxisome membrane acts as a sink for RNF146/TNKS, keeping RNF146/TNKS away from other substrates such as AXIN1 and Golgi-localized BLZF1, and thereby stabilizing them. In this model, the absence of peroxisomes allows RNF146/TNKS to re-localize to induce the degradation of AXIN1 and BLZF1. Indeed, reports in the literature suggest that both RNF146 and TNKS can re-localize in response to perturbations; RNF146 moves between the cytoplasm and nucleus in response to oxidative stress and DNA damage [Gero et al 2014; Kang et al 2011] and TNKS’s diffuse cytosolic localization becomes punctate with treatment with TNKS inhibitors [Martino-Echarri et al 2016, Thorvaldsen et al 2015] and infection with Sendai virus [Xu et al 2022].

Finally, we demonstrated that the effect of PEX knockdowns on the RNF146/TNKS substrate AXIN1 was sufficient to alter the transcriptional response to Wnt ligand in two different cell lines. Our results suggest that peroxisomes may act as signaling platforms that can alter cell fate decisions by impacting Wnt signaling. The most severe forms of Zellweger syndrome have stereotyped neuronal migration disorders, chondrodysplasia punctata, renal cysts, and craniofacial dysmorphisms indicating disruptions to normal development [Braverman et al 2016]. Our findings raise the possibility that the perturbation of cell signaling pathways contributes to the pathology of Zellweger Spectrum Disorders.

## Acknowledgements

We thank Katelynn Kazane for assistance with engineering CRISPRi cell lines. We thank Amos Liang and Jacob Corn for Hct116 CRISPRi cells. We thank Diego Acosta-Alvear for H4 CRISPRi cells. We thank Mary West and Pingping He of the High-Throughput Screening Facility (HTSF) at UC Berkeley for preparation of the pooled CRISPRi library and Jonathan Weissman for the gift of the pooled library. We thank Griffin Kramer with assistance in data processing. We thank members of the Gardner and Richardson labs, Joel Rothman, and Chris Hayes for fruitful discussions. We thank Ron Prywes and Randall Moon for plasmid reagents. Brooke Gardner acknowledges support from K99/R00GM121880, R35GM146784, and the Searle Scholars Program. Jonathan Vu acknowledges support from the Connie Frank Fellowship. Lori Mandjikian acknowledges support from CIRM Educ4. Chris Richardson acknowledges support from R35 GM142975. The content is solely the responsibility of the authors and does not necessarily represent the official views of the National Institutes of Health.

## List of Supplementary Figures

Figures S1 to S5

Table S1

## Materials and Methods

### Cell Lines, Culture Conditions, Lentiviral Production and Transduction

H4 dCas9-KRAB (a gift from the laboratory of Diego Acosta-Alvear, UCSB), HEK293T, and HCT116 dCas9-KRAB (a gift from the laboratory of J. Corn, ETH Zürich) cells were cultured in Dulbecco’s Modified Eagle Media (10565018, DMEM, Gibco) supplemented with 10% fetal bovine serum (FBS, S11150H, R&D Systems), 1% penicillin/streptomycin (15140122, Gibco), and 2 mM L-Glutamine, and kept at 37°C and 5% CO_2_ in a humidified incubator. Generation of lentivirus was performed by transfecting HEK293T cells with standard delta VPR and VSVG packaging vectors paired with TransIT-LTI Transfection Reagent (MIR2305, Mirus). Lentivirus was harvested 72 hrs following transfection and frozen at -80°C.

Zeocin resistance harboring HCT116 CRISPRi cell lines were constructed by transducing cells with lentivirus expressing mVenus-ZeoR-PTS1 constructs with either a PGK (pCR2054) or hEF-1α (pCR2055) promoter and ‘spinfecting’ cells in a centrifuge at 1000 rpm for 2 hrs. ZeoR expressing cells were single-cell sorted by flow cytometry (Sony SH800S) for mVenus expression at 488 nm excitation, where modestly fluorescent monoclonal cells were selected for both promoter types.

Re-expression constructs were made by Gibson cloning desired CDS sequences into the pLentiX-CD90 Thy1.1 vector backbone, with subsequent cell sorting of Thy1.1 positive cells by immunolabeling with CD90.1 Thy-1.1 antibody (17-0900-82, Thermo Scientific).

For drug treatment conditions, cells were treated with 50 nM bafilomycin (B1793, Sigma-Aldrich) for 15 hours, 5-10 μM hydroxychloroquine (H0915, Sigma-Aldrich) for 24 hours, 500 nM G007-LK (S7239, Selleck) for 24 hours, 10 μM XAV939 (575545, EMD Millipore) for 24 hours.

For siRNA treatment conditions, cells were transfected with 10 nM of desired siRNA using Lipofectamine RNAiMAX (13778150, ThermoFisher) according to the manufacturer’s protocol for 48 hrs.

### iPS cells

AICS-0090-391 (WTC-CLYBL-dCas9-TagBFP-KRAB-cl391) cells were cultured in 10 ml sterile-filtered mTeSR-Plus (100-0276, STEMCELL) on a Matrigel-coated plate (354277, Corning) and grown to 80% confluency, five days post-thaw at 37°C and 5% CO2 in a humidified incubator. For routine passaging, at 80% confluency, media was aspirated and cells were washed with 4 ml room temp DPBS prior to dissociation. iPSCs were then treated with 2 ml pre-warmed Accutase (AT104, Stem Cell Technologies) and the vessel was incubated at 37ºC for 10 mins. Once cells began to detach, 4 mLs DMEM/F12 were added to the Accutase-treated cells and dissociated cells were triturated. Cells were rinsed with an additional 7 ml of DPBS for a final wash, and the dissociated cell suspension was transferred to a 15 ml conical tube and centrifuged at 500g for 5 min at room temp. DMEM/F12/Accutase supernatant was carefully aspirated and cells were resuspended in 10 ml fresh mTeSR-Plus containing 10μM Y-27632 2HCl (ROCK Inhibitor, S1049, Selleck) (ROCKi) and counted via flow cytometry. Cells were then seeded into a Matrigel-coated six-well dish at a density of 1.5e+05 per well in 3 ml mTeSR-Plus containing ROCKi. Old media containing ROCKi was aspirated from each well the next day and replaced with fresh mTeSR1 without ROCKi. mTeSR-Plus was changed daily, and ROCKi was used for each passaging event, and always removed 24 hours thereafter.

### Genome-wide Pooled CRISPRi Screen and Analysis

HCT116 CRISPRi pCR2054 cells were transduced with lentivirus harboring constructs expressing sgRNAs from the genome-wide pooled CRISPRi v2 library with 8μg/mL of polybrene (TR-1003-G, EMD Millipore) at a multiplicity of infection (MOI) of <1. hCRISPRi-v2 library was a gift from Jonathan Weissman (Addgene ID #83969). Cells were then selected with 1.5 μg/mL puromycin (A1113803, Gibco) for 1 week in 15 cm dishes and expanded to 3.60 × 10^8^ cells to allow for T0 condition takedowns as well as base seed for Zeocin (R25001, Invitrogen) treated and untreated samples. Treated cells were subjected to Zeocin 25 ng/μL final concentration and untreated cells were substituted with DMSO. Cells were maintained at >500X coverage per library element per replicate per condition throughout the screen. Cells were then cultured for 35 days in 5-Chamber CellStack vessels, splitting cells every 48-72hrs. and harvesting 2.40 × 10^8^ cells every 7 days per condition, where the treated and untreated conditions reached ∼8 and ∼16 doublings at day 14, respectively. Genomic DNA was purified using Macherey-Nagel NucleoSpin Blood XL Maxi Kit (740950.50, Macherey-Nagel) and prepared as previously described [Kampmann et al 2014] with modifications: Sbf1 (R3642S, NEB) was used instead of PvuII for the restriction digest. Next-Generation Sequencing (NGS) was performed using an Illumina NovaSeq SP with 2×50 paired end reads using custom read primers:

Read 1:

GTGTGTTTTGAGACTATAAGTATCCCTTGGAGAACCACCTTGTTGG

Read 2:

CTAGCCTTATTTAAACTTGCTATGCTGTTTCCAGCTTAGCTCTTAAAC

NGS data was then quantified and phenotype scores were generated using python scripts from the Horlbeck Lab’s ScreenProcessing pipeline as previously described [Horlbeck et al 2016].

### Immunofluorescence Staining

Cells were plated on glass bottom 96-well plates and fixed using 4% paraformaldehyde (15710, Electron Microscopy Sciences) in DPBS (14190250, Gibco) for 10 minutes and washed twice with DPBS. Cells were then permeabilized using 0.25% Triton X-100 (A16046.AP, Thermo Fisher) in DPBS for 10 minutes, blocked with 3% BSA (BP9703100, Fisher Scientific) in PBST (DPBS, Gibco; 0.1% Tween 20, AAJ20605AP, Thermo Fisher) for 30 minutes, and then probed with desired antibody in 3% BSA PBST for 1 hour at RT. Cells were then washed 3 times with PBST and incubated with secondary antibody and DAPI (D1306, Invitrogen) in 3% BSA PBST for 1 hour at RT in darkness. Cells were then washed 3 times in PBST and stored in DPBS prior to image acquisition.

A list of antibodies used in the manuscript is available in **Table S1**.

### Confocal Microscopy and Analysis

Fluorescent image acquisition was performed using a Nikon Eclipse Ti2 configured with a spinning disk confocal scanner (Yokogawa, CSU-W1), CFI Plan Apochromat Lambda D 40X air objective lens, CFI Apochromat TIRF 100X/1.49 oil-immersion objective lens, and NIS-Elements AR software (Nikon, version 5.31.01). Green (mVenus), blue (BFP, DAPI), red, and far red were excited with 488, 405, 561, and 640 nm lasers, respectively. Microscopy images were post-processed using ImageJ/FIJI software (version 2.0.0). Quantification and analysis of microscopy images was performed using CellProfiler [Stirling et al 2021] (version 4.2.4). For live cell images, acquired images were thresholded by global minimum cross entropy to select for and differentiate between cell cytoplasm area and mVenus foci area in an unbiased manner; downstream mVenus foci number, area, and intensity was measured within a range of size and ROI. For immunofluorescence microscopy, images were processed by, first, defining nuclei stained by DAPI by adaptive Otsu 3-class thresholding to differentiate between background and nuclei; second, by expanding from nuclei objects to define cytoplasm based on distance and Otsu 2-class thresholding and then subtracting nuclei from this area; third, by selecting, within the cytoplasm area, foci objects for mVenus, Catalase, or PMP70 of a defined size and ROI determined by adaptive Otsu 3-class thresholding. All of the previously mentioned objects are then measured for number, area, intensity, and colocalization by Pearson’s correlation.

### Fluorescence-activated Cell Sorting

Flow cytometry was performed using an Attune NxT Flow Cytometer (Invitrogen) or SH800S (Sony). Excitation wavelengths of 488 nm (530/30 filter) and 405 nm (450/40 filter) were used to analyze mVenus and BFP expression, respectively. For selection of cells re-expressing PEX14 or PEX19, cells were sorted for Thy1.1 positive cells after immunolabeling with APC-conjugated CD90.1 Thy-1.1 antibody (17-0900-82, Thermo Scientific) at excitation wavelength of 638 nm (720/60 filter). FCS data was analyzed and visualized using FlowJo (version 10.6.2).

### Immunoblotting

Cells were trypsinized (0.05% Trypsin, 25300062, Gibco), quenched, spun down at 300 x g for 5 minutes, decanted, and washed using DPBS (14190250, Gibco). Cells were lysed using RIPA lysis buffer (0.1% SDS, BP8200100, Fisher Scientific, 1% IPEGAL CA630, 8896, EMD-Millipore; 0.5% sodium deoxycholate, D6750, Sigma, 50mM Tris, BP152-5, Fisher Scientific; 150mM NaCl, S271-10, Fisher Scientific) with benzonase (101697, EMD Millipore) and protease inhibitor (78430, Thermo Scientific) for 30 minutes on ice and spun down at 14,000 rpm for 5 minutes and supernatant collected. Total protein concentrations were quantified using Bio-Rad Protein Assay (5000006, Bio-Rad). Protein samples were normalized to 10-20 μg, mixed with 4X Laemmli sample buffer (62.5mM Tris, 10% glycerol, 1%SDS, 0.005% bromophenol blue) containing beta-mercaptoethanol (M6250, Sigma-Aldrich), and incubated for 5 minutes at 95 deg C. Samples were loaded and resolved on 4-20% SDS-PAGE gels (#4561095, Bio-Rad), semi-dry transferred to 0.45 μm LF PVDF membranes (1620264, Bio-Rad), blocked in 5% milk (Nestle) in TBST (50mM Tris, 150mM NaCl, pH 7.4), and probed with desired antibody in 3% BSA TBST (BSA, BP9703100, Fisher Scientific) overnight at 4C. Membranes were then washed and probed with secondary HRP-conjugated antibodies, with visualization of chemiluminescence using Pierce ECL2 Western Blotting Substrate (PI80196, Thermo Scientific) on a ChemiDoc MP Imaging System (Bio-Rad).

### TOPFlash

HEK293T ZIM3-dCas9 and HCT116 dCas9-KRAB cells harboring NTC, PEX14, and PEX19 sgRNAs were transfected in 96-well plates by lipofectamine (TransIT LT-1, Mirus). A normalized 55 ng of total plasmid DNA was used at a ratio of 50:5 TOPFlash/FOPFlash:Renilla. Cells were treated with either BSA or human recombinant WNT3a (5036-WN-010, R&D Systems) 24 hours later. Cells were then lysed 24 hours after treatment and luciferase activity was measured using the Dual Luciferase Assay (E1910, Promega) according to manufacturer’s protocol, and luminosity was read out using a microplate reader (SpectraMax M5, Molecular Devices). Plasmids used were: TopFLASH (Addgene#12456), FopFLASH (Addgene#12457), Renilla (Addgene#27163). M50 Super 8x TOPFlash and M51 Super 8x FOPFlash (TOPFlash mutant) were a gift from Randall Moon (Addgene plasmid # 12456, #12457) [Veeman et al 2003]. pRL-SV40P was a gift from Ron Prywes (Addgene plasmid # 27163) [Chen and Prywes 1999].

### Crude Fractionation

Designated cell lines were harvested at 10-15 million cells (equalized among experimental replicates), spun down, and resuspended in Homogenization Buffer (250mM Sucrose, 20mM HEPES, 10mM KCl, 1.5mM MgCl2). Cells were quickly freeze thawed and mechanically homogenized via dounce, with a minimum of 10 passes, to lyse the extracellular membrane while retaining intracellular organelles. Total was collected, and the remainder of the product was fractionated at 20,000xg for 15 minutes at 4°C (C1). The fractionated supernatant was harvested, leaving behind the organellar pellet, and then further purified by centrifugation at 20,000xg for 15 minutes at 4°C (C2) again to remove any remaining membranes. The resulting supernatant is considered the cytoplasmic fraction. The pellet from the primary centrifugation (C1) is washed by resuspension in 1 mL of homogenization buffer, spun down at 20,000xg for 15 minutes at 4°C (C3), the wash supernatant is discarded, and the pellet is lysed by RIPA lysis buffer; this is considered the cell pellet.

### Co-Immunoprecipitation

Designated cell lines were harvested at 10-15 million cells (equalized among experimental replicates), spun down, and resuspended in LB1 (60 mM HEPES pH 7.6, 150 mM NaCl, 150 mM KCl, 10 mM MgCl2, 0.2% IPEGAL CA630 (Sigma), 0.1% sodium deoxycholate (Sigma), 1X Protease Inhibitor, 1X Benzonase). The suspension quickly undergoes freeze-thaw, is then dounce homogenized (with a minimum of 10 passes), and then is incubated for 30 minutes at 4°C with inversion. Total lysate samples are acquired and the remaining lysate is centrifuged for 5 min, 20,000xg at 4°C to separate the sample into soluble and insoluble fractions. The supernatant (soluble fraction) is collected and spun at 20,000 xg at 4°C for 30 minutes to clear out any remaining insoluble proteins or cell debris, this is the lysate supernatant. The insoluble fraction is washed with LB1, spun down, decanted, and resuspended in LB1, this is the pellet. The lysate supernatant is pre-cleared with protein G agarose beads for 30 minutes at 4°C and washed. For FLAG-IPs, supernatant is incubated with M2 FLAG conjugated agarose beads (M8823, Millipore) for 3 hrs at 4°C with inversion. For TNKS immunoprecipitations, supernatant is incubated with 10 μg of TNKS antibody (sc-365897, Santa Cruz Biotech) for 3 hrs at 4°C with inversion, and then conjugated to Protein G Dynabeads (10004D, Invitrogen) for 3hrs. at 4°C with inversion. Beads are then washed 5X in LB2 buffer (LB1 buffer without protease inhibitors or benzonase) with either centrifugation or magnetic stand (where applicable). Beads are then eluted using 50 μL of freshly prepared (day of) Elution Buffer (100 mM NaHCO3, 1% SDS) at 65°C for 15 min on a heated shaker (1200 rpm) twice.

### RNA-Seq

HCT116 Pex-ZeoR cell lines harboring either NTC or RNF146 sgRNAs were harvested, spun down, and RNA was extracted using RNeasy Mini Kit (Qiagen #74104) according to manufacturer instructions, in triplicate. Purified RNA samples were poly-(A) enriched, reverse transcribed, and sequenced on an Illumina NovaSeq to produce paired-end 150 bp reads (Novogene). Raw fastq reads were trimmed using fastp v0.23.2 [Chen et al. 2018], alignment was done via STAR v 2.7.11a [Dobin et al. 2013], count tables were generated using htseq2 v. 2.0.2 [Givanna et al. 2022] and differential expression analysis was performed using the R-package DESeq2 v. 1.40.1 [Love et al. 2014]. Differential expression comparisons were made between experimental and nontargeting CRISPRi strains in biological triplicate.

## Notes

### Competing Interest Statement

The authors have declared no competing interest.

