## Supplementary Figures for "A genome-wide screen links peroxisome regulation with Wnt signaling through RNF146 and tankyrase"

* co-corresponding

#### Supplementary Figures

**Figure S1. (A)** Quantification of cell count by flow cytometry in different concentrations of Zeocin of HCT116 cells with sgRNAs targeting NTC, PEX1, or PEX6, over 72 hrs. Data is representative of n=2 biological replicates. Cell count is normalized to untreated. **(B)** Quantification of flow cytometry data of BFP- (NTC) and BFP+ (PEX6) cells grown in co-culture competition assay over t=11 days in the presence of 0, 25, or 50 ng/uL of Zeocin. Timepoints are taken every t=2 days. Data shown as the mean ± SD of n=3 biological replicates. **(C)** Schematic of the CRISPRi screen. Pex-ZeoR cells were transformed with a genome-wide gRNA library, selected for expression of guides, and split into untreated and +Zeocin growth conditions. Genomic DNA takedowns for NGS sequencing at t=0 and t=7x for all conditions. **(D)** Heatmap showing Pearson's correlation coefficient of guide abundance for all library elements between biological replicates of sequenced timepoints between treated and untreated conditions. T and Z represent untreated and Zeocin treated conditions, respectively, while numbers represent timepoint (days). Highlighting indicates comparisons between day 14 samples. **(E)** Fold change of various PEX sgRNA abundances derived from genome-wide CRISPRi screen comparing Zeocin treated to untreated samples. Highlighting indicates comparisons between day 14 samples. Y-axis is phenotype score, a measure of fold change of 3 of 5 significant guides per gene. X-axis is time (t) in days. Data is representative of n=2 biological samples.

### Supplementary Figure 1

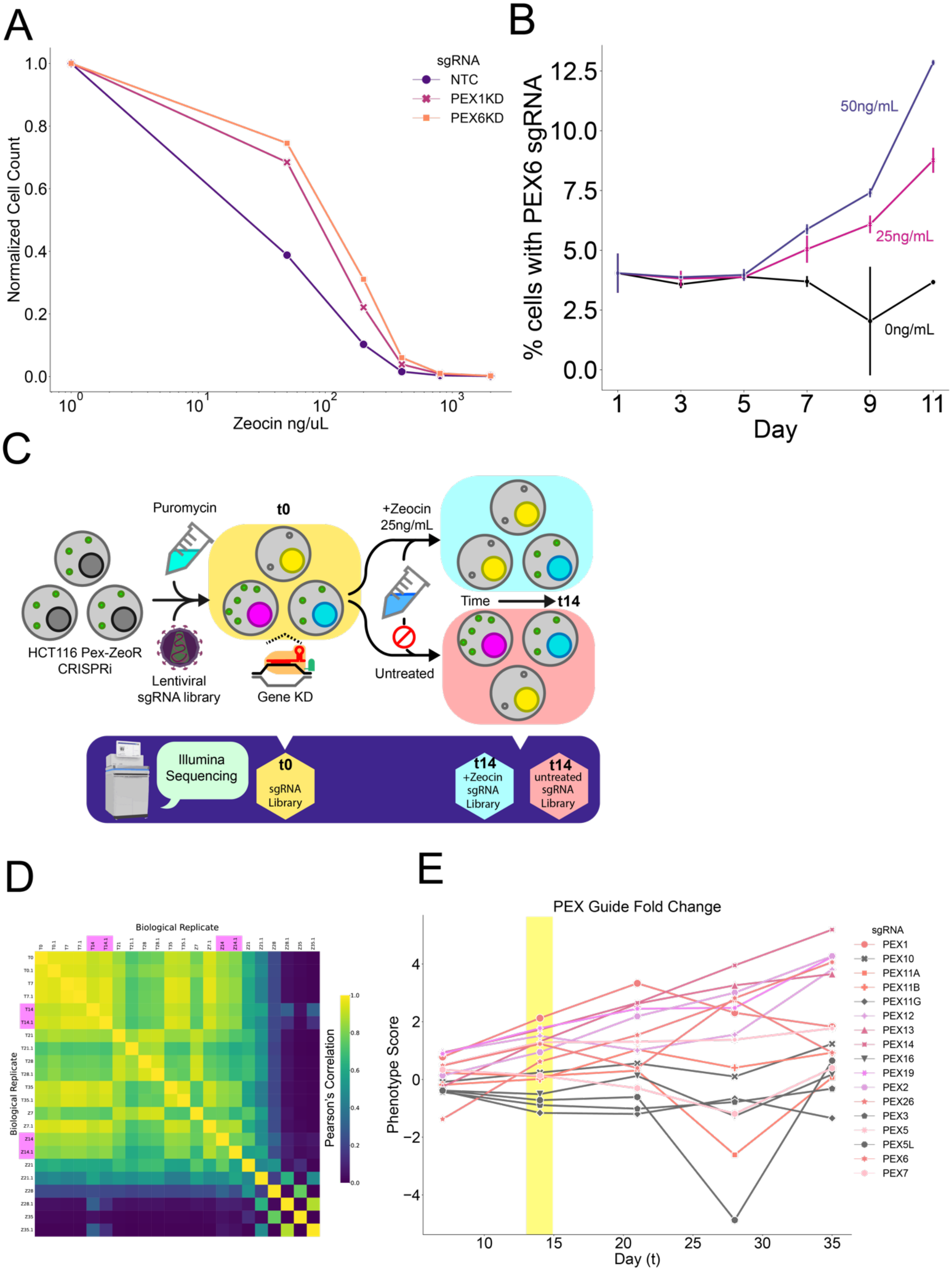

**Supplementary Figure 2. (A)(B)** Additional data as in Figure 2A CellProfiler quantification of the ratio of mVenus-PTS1 intensity in foci and in the cytoplasm in fluorescence microscopy images acquired of live HCT116 Pex-ZeoR cells expressing sgRNAs targeting various genes. (A) Positive phenotype score genes from the primary genetic screen. (B) Negative phenotype score genes from the primary genetic screen. Data per gene constitutes m=2 unique sgRNAs with n=49 images per gene. Non-targeting control sgRNA shown in yellow, PEX1 sgRNA shown in pink, sgRNAs significantly different from NTC are in blue (p<0.0001, independent t-test) or purple (p<0.05, independent t-test), and sgRNAs with p>0.05 are in white.

#### Supplementary Figure 2

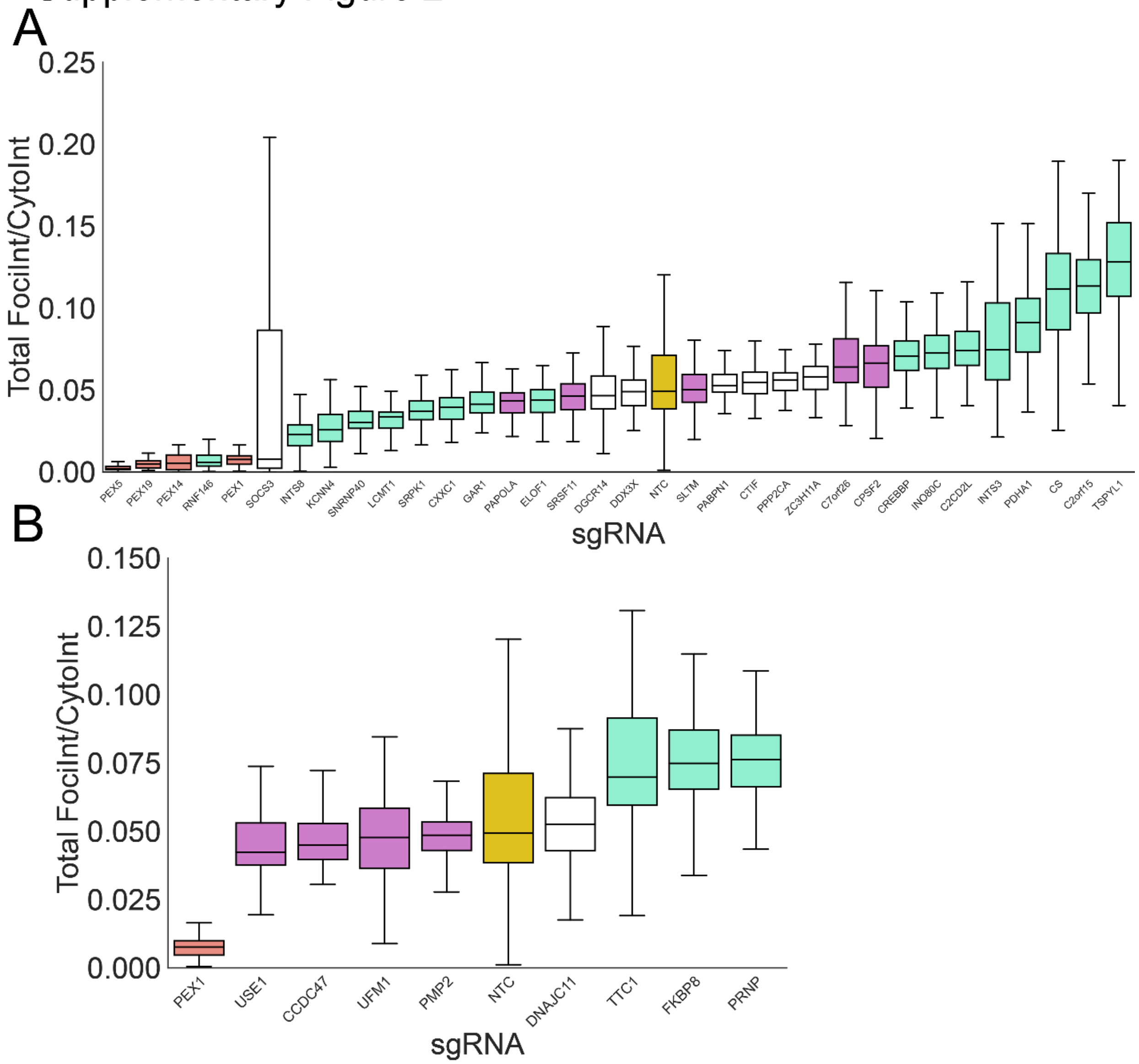

**Figure S3. (A)** Left Panel: Representative immunofluorescence microscopy images of NTC and RNF146 sgRNA expressing HCT116 Pex-ZeoR cells treated with DMSO (mock) or 50 nM Bafilomycin A1 (Baf) for 15 hrs. mVenus-PTS1 in green, DAPI in blue, and PMP70 in cyan. Scale bar: 10 µm. Right panel: Immunoblot of TNKS and LC3B of cell lysate from conditions in left panel. **(B)** Quantification of immunofluorescence microscopy images in A for percentage foci area of mVenus-PTS1 and PMP70 versus cytosolic area for m=21 images and n=2 biological replicates. **(C)** Left Panel: Representative immunofluorescence microscopy images of NTC and RNF146 sgRNA expressing HCT116 Pex-ZeoR cells treated with DMSO (mock), 5 uM hydroxychloroquine (HCQ), or 10 uM hydroxychloroquine for 24hrs (5uM HCQ not shown). mVenus-PTS1 in green, PMP70 in magenta, and DAPI in blue. Scale bar: 10µm. Right panels: Quantification of immunofluorescence microscopy images for percentage foci area of mVenus and PMP70, respectively, versus cytosolic area. for m=32 images and n=2 biological replicates. **(D)** Immunoblots of cellular lysate from (C) against TNKS and LC3B. **(E)** Left Panel: Representative fluorescence microscopy images of NTC and RNF146 sgRNA expressing H4 Pex-ZeoR cells treated with DMSO (mock) or 50 nM Bafilomycin A1 for 15 hrs. Scale bar: 10 µm. Middle panel: Quantification of mVenus-PTS1 microscopy images in left panel for mVenus foci intensity (peroxisomes) versus total cytosol intensity. Data is representative of m=32 images per condition and n=2 biological replicates. Right Panel: Immunoblots of cellular lysate from left panel against TNKS and LC3B. Data shown are representative of n=3 independent blots. Asterisks denote p-values *p <0.05, **p <0.01, ***p <0.001, ****p <0.0001, whereas ns denotes not significant, calculated by independent t-test.

### Supplementary Figure 3

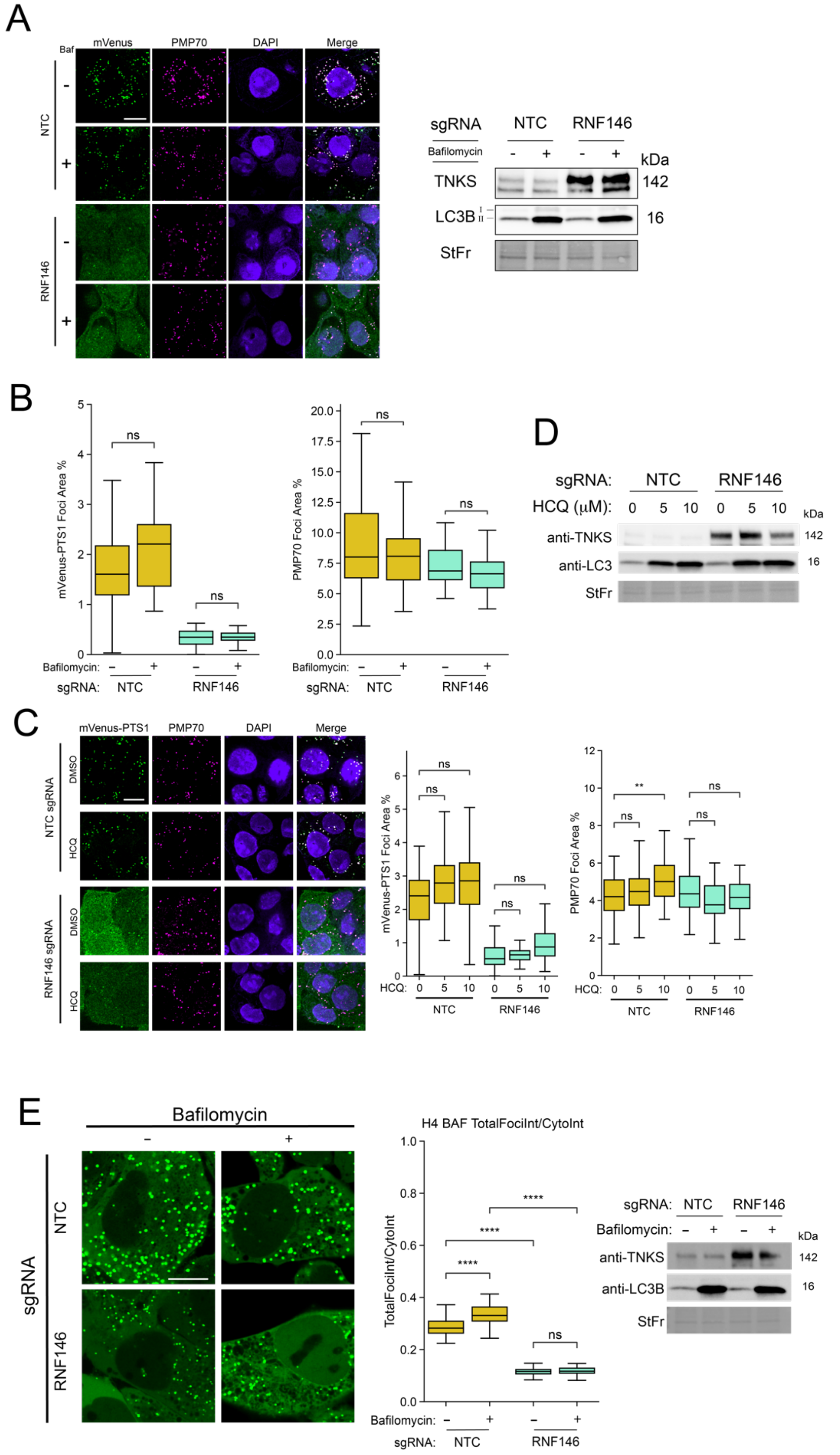

**Figure S4. (A)** Left Panel: Representative immunofluorescence microscopy images of NTC, RNF146, PEX19, and PEX5 sgRNA expressing H4 Pex-ZeoR cells. mVenus-PTS1 in green, DAPI in blue, PMP70 in magenta. **(B, C)** Quantification of immunofluorescence microscopy images for percentage foci area of PMP70 **(B)** and mVenus **(C)**, respectively, versus cytosolic area. n=25 images. **(D)** Left Panel: Representative immunofluorescence microscopy images NTC, RNF146, PEX19, and PEX5 sgRNA expressing cells. Catalase in yellow, DAPI in blue, PMP70 in magenta. m=25 images. n=2 biological replicates. Right panels: Quantification of Pearson's correlation coefficient of catalase and PMP70 colocalization of microscopy images.

### Supplementary Figure 4

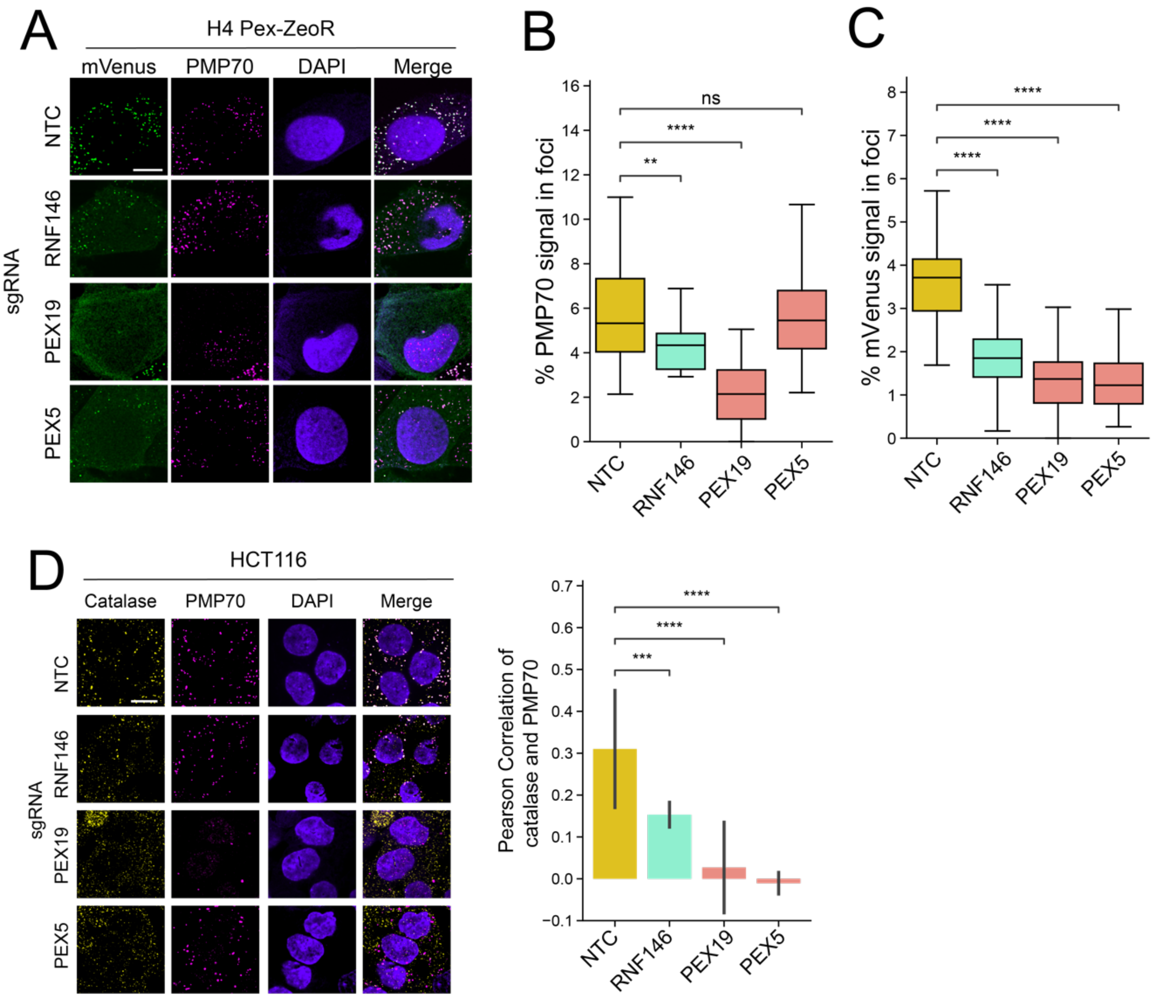

**Figure S5. (A)** Immunoblot measuring the abundance of AXIN1 in H4 CRISPRi cells expressing sgRNA for NTC, PEX5, PEX14, PEX19, and RNF146. Blots are representative of n=3 blots.

### Supplementary Figure 5

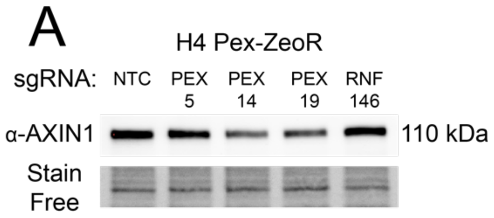

**Table S1. List of Antibodies**

| Target | Manufacturer | Catalog Number |
|---|---|---|
| ACTB | Sigma | A2228 |
| ATG7 | Cell Signaling Technologies | 2631T |
| AXIN1 | Invitrogen | MA5-14853 |
| BLZF1 | ThermoFisher | MA5-27126 |
| CASC3 | Proteintech | 18047-1-AP |
| Catalase | Invitrogen | LF-MA0004 |
| CD90 | Thermo Scientific | 17-0900-82 |
| FLAG | Sigma | F1804-200UG |
| GAPDH | ThermoFisher | MA5-15738 |
| LC3 | Cell Signaling Technologies | 2775 |
| PEX14 | Proteintech | 10594-1-AP |
| PEX5 | ThermoFisher | PA5-58717 |
| PMP70 | Invitrogen | PA-650 |
| TNKS | Santa Cruz Biotechnology | sc-376875 |
